# Flexibility and rigidity index for chromosome packing, flexibility and dynamics analysis

**DOI:** 10.1101/374132

**Authors:** Jiajie Peng, Jinjin Yang, Kelin Xia

## Abstract

**Motivation:** The packing of genomic DNA from double string into highly-order hierarchial assemblies has great impact on chromosome flexibility, dynamics and functions. The open and accessible regions of chromosome are the primary binding positions for regulatory elements and are crucial to nuclear processes and biological functions.

**Results:** Motivated by the success of flexibility-rigidity index (FRI) in biomolecular flexibility analysis and drug design, we propose a FRI based model for quantitatively characterizing the chromosome flexibility. Based on the Hi-C data, a flexibility index for each locus can be evaluated. Physically, the flexibility is tightly related to the packing density. Highly compacted regions are usually more rigid, while loosely packed regions are more flexible. Indeed, a strong correlation is found between our flexibility index and DNase and ATAC values, which are measurements for chromosome accessibility. Recently, Gaussian network model (GNM) is applied to analyze the chromosome accessibility and a mobility profile has been proposed to characterize the chromosome flexibility. Compared with GNM, our FRI is slightly more accurate (1% to 2% increase) and significantly more efficient in both computational time and costs. For a 5kb resolution Hi-C data, the flexibility evaluation process only takes FRI a few minutes on a single-core processor. In contrast, GNM requires 1.5 hours on 10 CPUs. Moreover, interchromosome information can be easily incorporated into the flexibility evaluation, thus further enhance the accuracy of our FRI. In contrast, the consideration of interchromosome information into GNM will significantly increase the size of its Laplacian matrix, thus computationally extremely challenging for the current GNM.

**Availability:** The software is available at https://github.com/jiajiepeng/FRI_chrFle.

## 1 Introduction

The packing of chromosome into complicated three-dimensional hierarchial structure has a profound effect on gene expression and other biological functions (Schmitt *et al.*, 2016a). For instance, chromatin loop is formed when cis-regulatory element, such as enhancers, are folded into close spatial proximity with its target promoter. This long-range chromatin contacts are vital to the regulation of gene expression. Recently, the 4D nucleome project is proposed to reveal the packing and dynamics of chromosome and gain insight on the mechanism of gene regulation (Dekker *et al.*, 2017). A major driving force for the project is the advancement of genome-wide C-techniques and C-data for multiple species and tissues (Dekker *et al.*, 2002; Simonis *et al.*, 2006; Zhao *et al.*, 2006; Fullwood *et al.*, 2009; de Wit and de Laat, 2012; Lieberman-Aiden *et al.*, 2009; Dixon *et al.*, 2012; Nora *et al.*, 2012; Jin *et al.*, 2013; Bonev and Cavalli, 2016; Schmitt *et al.*, 2016b; Nagano *et al.*, 2013). With the structure information obtained from C-techniques, researchers begin to understand more about packing and organization principles of chromosomes. In general, the structure of mammalian chromosomes can be explored from several different scales (Bonev and Cavalli, 2016), including nucleosome, chromatin fiber, chromatin loops (Rao *et al.*, 2014a), topological associated domain (TAD) (Dixon *et al.*, 2012; Nora *et al.*, 2012), genomic compartment (Lieberman-Aiden *et al.*, 2009), chromosome territory (Lieberman-Aiden *et al.*, 2009), etc. A nucleosome is a basic building block for chromatin organization. They interact with each other to form the 30 nm chromatin fibres with solenoid or zigzag shape. These chromatin fibres aggregate and form chromatin loops, in which cis-regulatory element, such as enhancers, are folded into close spatial proximity with its target promoter. From Hi-C data analysis, larger-scale structures, i.e., TAD and genomic compartment, have been defined. TADs, which are about 200 kilobases(Kb) to 2 megabases(Mb), are chromosome components that are highly consistent between different cell types and species. Genomic compartment, which is classified into type A and type B, represents chromosome regions that either densely or sparsely packed. The packing of chromosome into its hierarchial structure is greatly facilitated by various insulator proteins, cohesin complex, mediator, border elements, loop-extruding complexes, other DNA-binding proteins, as well as RNAs. Algorithms and models that based on C-data and ENCODE data are proposed (Lieberman-Aiden *et al.*, 2009; Dixon *et al.*, 2012; Filippova *et al.*, 2014; Lévy-Leduc *et al.*, 2014; Baù *et al.*, 2011; Hu *et al.*, 2013; Zhang *et al.*, 2013; Segal *et al.*, 2014; Lesne *et al.*, 2014; Zhang and Wolynes, 2015; Imakaev *et al.*, 2015). However, even with these progresses, there is a lacking of physical models that quantitatively analyzes the chromosome packing, flexibility and dynamic properties.

In structure biology, it is well-known that biomolecular flexibility, dynamics and functions are tightly related to their structures. For instance, in an intrinsically disordered protein, its well-organized regions are usually very rigid and highly stable. In contrast, its disorder parts, such as hanging chains and extruding loops, are normally very flexible and easy to interact with others. In fact, flexible regions in biomolecules are always more dynamic and tend to have interactions with ligands or other biomolecules. Experimentally, flexility can be quantitatively measured in terms of Debye-Waller factor (or B-factor). Various models have been proposed to reveal the deep connection between biomolecular structure and flexibility (Flores *et al.*, 2007; Emekli *et al.*, 2008; Keating *et al.*, 2009; Shatsky *et al.*, 2004; Flores and Gerstein, 2007; Tama *et al.*, 2000; Halle, 2002; Kundu *et al.*, 2002; Kondrashov *et al.*, 2007; Song and Jernigan, 2007; Hinsen, 2008; Park *et al.*, 2013; Demerdash and Mitchell, 2012; Zhang and Brüschweiler, 2002; Lin *et al.*, 2008; Huang *et al.*, 2008; Li and Brüschweiler, 2009). Among these methods, flexibility-rigidity index (FRI) (Xia *et al.*, 2013; Opron *et al.*, 2014) is one of the most accurate and efficient model in B-factor prediction. In FRI model, a biomolecular structure is viewed as an equilibrium state in which all the interactions from the surrounding environment and within the molecule are well balanced. The unique position of each atom in the structure is the outcome from the “fight" with all the other atoms. Therefore, instead of resorting to the complicated protein interaction Hamiltonian as in other methods, FRI measures the biomolecular flexibility by its topological connectivity or packing density (Halle, 2002).

Compared with the classic models, such as Gaussian network model (GNM) and anisotropic network model (ANM), FRI has significantly increased the accuracy and dramatically reduced the computational time. In both GNM and ANM, a large matrix, either Laplacian matrix or Hessian matrix, is constructed. Their B-factor prediction formula requires the calculation of all the eigenvalues and eigenvectors of the matrix, which can be time-consuming not to mention about the memory cost. Free from eigenvalue decomposition, FRI only has the computational complexity of *𝒪*(*N*^2^) with *N* the total number of atoms. Fast FRI (fFRI) (Opron *et al.*, 2014) can further reduce the computational complexity to *𝒪* (*N*) with almost no scarifying the accuracy of the model. It only takes fFRI 30 seconds on a single-core processor to calculate the flexibility of an HIV virus capsid with 313 236 residues (Opron *et al.*, 2014). Multiscale FRI (Opron *et al.*, 2015a), Generalized FRI (Nguyen *et al.*, 2016) and multiscale weighted colored graph (MWCG) based FRI (Bramer and Wei, 2018) can further increase the accuracy of FRI models. Especially, MWCG based FRI has set a new accuracy benchmark for protein flexibility analysis. More interestingly, FRI model has been applied to protein-ligand binding affinity prediction in drug design (Nguyen *et al.*, 2017). The results from the FRI based machine learning model has significantly outperformed all traditional models. This reveals that the rigidity strengthening can be a potential mechanism for protein-ligand binding (Nguyen *et al.*, 2017). More recently, a virtual particle based FRI model is proposed for analyzing the dynamics of extremely large-sized biomolecular complexes and organelles, especially the ones from Electron Microscopy Data Bank (EMDB) (Xia and Wei, 2016; Xia *et al.*, 2018). The success of the FRI model in these subcellular structures have motivated us to propose the FRI based model for chromosome packing, flexibility and dynamics analysis.

Recently, Gaussian network model has been used to quantitatively characterize the chromatin accessibility. It is known that packing of DNA into a compressed form causes accessibility problems for transcription and DNA replication. In general, the chromatin packing density is negatively correlated with transcriptional activity. For tightly packed chromatin regions, their DNA is less accessible to transcriptional machinery. For loosely packed chromatin regions, the DNA is more accessible and transcription is much easier. In this way, the chromatin accessibility can be characterized by nuclease hypersensitivity, which is directly measured by MNase, DNase, FAIRE, as well as ATAC. Based on the Hi-C data, a mobility profile is defined from GNM and has been found to be highly correlated with DNase-seq and ATAC-seq values (Sauerwald *et al.*, 2017). Essentially, the mobility profile can be viewed as a characterization of flexibility properties of the chromosome structures.

In the current paper, we propose a FRI based model for chromosome flexibility analysis. Similar to biomolecular flexibility, we assume the rigidity and flexibility of each loci is solely determined by its local packing density. In this way, the rigidity index for each locus can be directly evaluated by summarizing its connectivity with its local neighbours. The connectivity strength between any two loci can be evaluated from its contact frequency from the Hi-C matrix. The flexibility index is inversely related to the rigidity index and can be used to characterize the chromatin accessibility. Our flexibility model is validated by comparing with the chromosome packing information from the chromatin accessibility. A high correlation coefficient is found between our flexibility index, DNase-seq and ATAC-seq values. Compared with GNM (Sauerwald *et al.*, 2017), our model is slightly more accurate with average 1%-2% increase in accuracy and significantly more efficient in both computational time and resources. Moreover, interchromosome interaction can be incorporated into our FRI model. As expected, the inclusion of the interchromosome interactions further enhances the performance of our model. Essentially, our FRI model provides a direct link between the chromosome packing density with chromosome flexibility.

## 2 Methods

### 2.1 Flexibility-rigidity index

#### Basic setting of flexibility-rigidity index

The flexibility and rigidity property of a systems is highly related to their inner network structure. We consider a system of *N*-elements with coordinates {**r**_*j*_|**r**_*j*_ ∈ ℝ^3^*, j* = 1, 2, …, *N*}. The connectivity between the *i*-th and *j*-th element can be characterized by a general correlation kernel Φ(∥**r**_*i*_ − **r**_*j*_∥; *η_ij_*) satisfying,

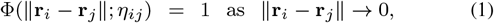

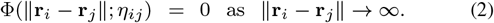

The parameter *η_ij_* is a resolution parameter (or scale parameter). And the correlation kernel can be chosen as any real-valued monotonically decreasing radial basis function. The commonly used kernel functions (Xia *et al.*, 2013; Opron *et al.*, 2014, 2015b) include generalized exponential functions

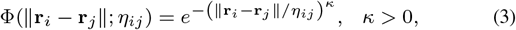

and generalized Lorentz functions,

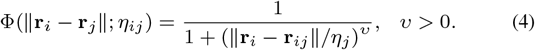

For *i*-th element, the rigidity index *μ_i_* is defined as the summation of its connectivity with all other atoms

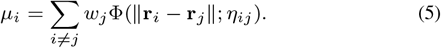

Here the weight parameter *w_j_* is used to characterize the atom properties, for example, it can be chosen as the atomic number. The flexility index is inversely related to the rigidity index. For *i*-th element, the flexility index *f_i_* is defined as,

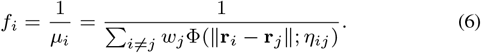

The flexibility index *f_i_* is linearly related to *i*-th B-factor,

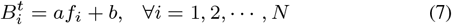

where 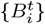 is the predicted B-factor. Linear regression model can be used to determine the fitting parameter *a* and *b*. In biomolecular flexibility analysis, a coarse-grained *Cα* representation is usually considered. In this case, we can set the weight parameter *w_i_* = 1*, i* = 1, 2*, …, N* and scale parameter *η_ij_* = *η, i, j* = 1, 2*, …, N* with *η* a fixed scale value. The fast FRI (fFRI) model is proposed for modeling extremely large biomolecules (Opron *et al.*, 2014). Essentially, a cell list algorithm (Allen and Tildesley, 1987) is employed and the rigidity index in Eq. (5) is calculated between atoms within a certain cut-off distance.

#### Rigidity function and flexibility function

Mathematically, flexibility and rigidity index are discrete values defined on atoms. They can be generalized into continuous representations, i.e., flexibility function and rigidity function (Xia *et al.*, 2013; Opron *et al.*, 2014). More specifically, the rigidity function can be defined as follows,

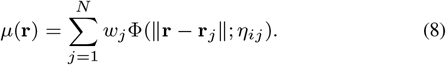

Essentially, a continuous rigidity function can be viewed as a density distribution and equals to Gaussian surface (Liu *et al.*, 2015) when *κ* = 2 and *w_i_* is chosen as the atomic number. A continuous flexibility function can be defined in a similar way as the flexibility index,

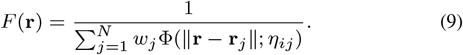

It should be noticed that the flexibility function is well defined only in the region when rigidity index is nonzero.

#### Multiscale flexibility-rigidity index

For a system with a hierarchial structure and multiscale properties, a unique scale parameter value is usually not suitable. Therefore, multiple correlation kernels with different scale values are considered. The multiscale flexibility (Opron *et al.*, 2015a) can be generalized as follows,

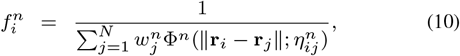

where 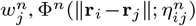 and 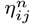 are quantities associated with *n*-th kernel. The linear regression is used to minimize the objective function,

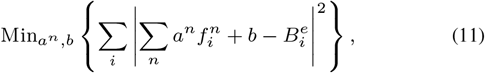

where 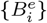 are the experimental B-factors. Usually, for each kernel function, a unique scale parameter *η* is used. The multiscale properties can be well characterized by several different *η* values.

#### Multiscale weighted colored graph based FRI

As a special case of graph labeling, graph coloring is to assign labels (or "colors") to nodes or edges of a graph under a certain rule or some constraints. In the weighted colored graph model, a protein graph is labelled by its edge types and subgraphs are defined according to these labels (Bramer and Wei, 2018). The atoms in a protein structure are predominately from several atom types, including carbon (C), nitrogen (N), oxygen (O), and sulfur (S). Hydrogen and ion atoms (such as Mn^2+^, Mg^2+^, Fe^2+^, Zn^2+^, etc) are not considered due to their absence from most PDB files. With this setting, all nodes in a colored protein graph can be labeled by element in an atom set (C, N, O, S) and all edges can be colored by element-specific pairs in a set (CC, CN, CO, CS, NC, NN, NO, NS, OC, ON, OO, OS, SC, SN, SO, SS). It should be noticed that edges in a colored protein graph are directed (Bramer and Wei, 2018). A CN pair is different from a NC pair in the colored graph model. A subgraph contains only the same type of directed pairs. For example, all NC pairs together form a subgraph. Further, these colored graphs are combined with FRI models, particularly the mFRI, to evaluate the protein B-factors. More specifically, if we are interested about flexility for all C atoms, we can consider subgraphs made from CC, CN, CO and CS pairs. For each subgraph, more than one scale values can be used. In this way, a multiscale weighted colored graph based FRI model is constructed.

### 2.2 FRI based chromosome packing density analysis

#### 2.2.1 Data processing

##### Hi-C data normalization

One of the major challenges for Hi-C data analysis is the systematic bias from the experimental setting that complicates the interpretation of observed contact frequencies (Yaffe and Tanay, 2011; Imakaev *et al.*, 2012; Ay *et al.*, 2014; Witten and Noble, 2012). Bias can be introduced from procedures including crosslinking, chromatin fragmentation, biotin-labelling and religation. Particularly, systematic biases that can substantially affect the Hi-C experimental results come from three major sources (Yaffe and Tanay, 2011), including distance-between restriction sites, the GC content and sequence mappability. Accounting for these biases is the first and most important step in C-data analysis. Various methods have been proposed to remove these biases, including HiCNorm (Hu *et al.*, 2012), Vanilla-Coverage normalization (Rao *et al.*, 2014a; Lieberman-Aiden *et al.*, 2009), iterative correction and eigenvector decomposition (ICE) (Imakaev *et al.*, 2012), Matrix-balancing method (Knight and Ruiz, 2013), etc. In this paper, we use the Vanilla-Coverage normalization.

##### Distance matrix construction

To construct the chromosome three-dimensional structure from the Hi-C data, computational models usually employ a reciprocal function to describe the relation between interaction frequency and spatial distance of two loci (Zhang *et al.*, 2013; Boulos *et al.*, 2013; Wang *et al.*, 2013; Segal *et al.*, 2014; Siahpirani *et al.*, 2016; Filippova *et al.*, 2014; Imakaev *et al.*, 2015; Lesne *et al.*, 2014; Chen *et al.*, 2016; Tjong *et al.*, 2016; Zhu *et al.*, 2018). More specifically, it assumes that the conversion between the contact frequency matrix *U* = {*u_ij_*; *i, j* = 1, 2*, …, N*} and the distance matrix *D* = {*d_ij_*; *i, j* = 1, 2*, …, N*} follows a power law distribution *d_ij_* = 1*/*(*u_ij_*)^*α*^. The coefficient *α* is a parameter called conversion factor, and *d_ij_* and *u_ij_* are the distance and contact frequency between loci i and j, respectively. The relation can be expressed as follows,

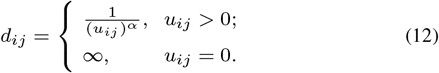

In this paper, we assume the conversion factor *α* = 1. Even though it is unphysical to assume the spatial distance equals to infinity when a contact frequency is zero, this definition works well for the kernel functions in our FRI models.

#### 2.2.2 Algorithm

We assume that the interaction between two loci in a chromosome structure can be characterized by the general correlation kernel as in Eqs. (1) and (2). We assume all loci have similar properties, thus *w_ij_* = 1 and *η_ij_* = *η*. The value of scale parameter *η* are linearly related to the locus resolution. Generally, a small *η* value is used in modeling high resolution data, whereas a large scale value in modeling low resolution ones. As shown in Algorithm 1, the process of chromosome flexibility analysis includes four components: normalizing the Hi-C data; transfering Hi-C contact frequency to relative distance; calculating the chromosome rigidity; calculating the chromosome flexibility.

**Algorithm 1.**
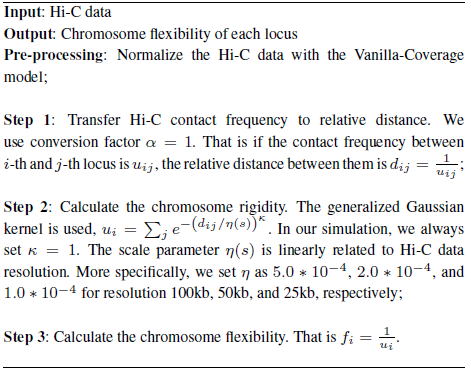
Chromosome flexibility analysis with FRI

## 3 Results and discussions

### 3.1 Data Description

To evaluate the performance of our FRI models, we consider the Hi-C dataset for GM12878 and IMR90 cell line with GEO accession number GSE63525 from Rao’s paper (Rao *et al.*, 2014b). GM12878 is a lymphoblastoid cell line produced from the blood. IMR90 is a cell line derived from human foetal lung. We also evaluate the Spearman correlation coefficient (SCC) between chromosome flexibility and chromosome accessibility measurements, including Dnase and ATAC. The DNase-seq data are obtained from ENCODE project. The ATAC-seq data is obtained from GEO database (GEO accessions GSM1155959 for GM12878 and GSM1418975 for IMR90). For both experimental datasets, bed-formatted peak files are used. We bin the data into the same resolution as used in the Hi-C data by adding all peak values within each locus. The binned data were then smoothed using moving average with a window size of 200 kb in the same way as GNM (Sauerwald *et al.*, 2017).

### 3.2 FRI based chromosome flexibility analysis

As stated in the introduction, there is a strong correlation between chromosome packing density, accessibility and flexibility. To illustrate their relations, we consider two 5kb resolution Hi-C chromosome data from flexibility index from our FRI. Chromatin accessibility is characterized by DNase-seq and ATAC-seq values (Sauerwald *et al.*, 2017). After the normalization of DNase-seq and ATAC-seq data, flexibility index is linearly fitted with their values. The results are demonstrated in Figure 1. For DNase-seq, it can be seen that there is a very good agreement between the predicted values and experimental ones. For ATAC-seq, the results are not as good as DNase-seq models. But a comparably good agreement is still observed. To further evaluate our model, we consider all the 23 chromosomes from GM12878 and IMR90, and calculate the Spearman correlation coefficient (SCC) between experimental results and theoretical predictions. The resolution of the Hi-C data is 25kb. Figures 2 and 3 demonstrate the SCCs for cell lines GM12878 and IMR90, respectively. The results for DNase-seq and ATAC-seq are listed in subfigures (*a*) and (*b*). In DNase-seq models, the average SCCs of all 23 chromosomes for GM12878 and IMR90 are 0.860 and 0.847 respectively. In ATAC-seq models, the average SCCs for GM12878 and IMR90 are 0.633 and 0.674. To avoid confusion, the parameter values of *κ* and *η* in our FRI model are set as 1.0 and 5.0 * 10^*−*4^ for GM12878, 1.0 and 3.0 * 10^*−*3^ for IMR90, respectively.

**Fig. 1.**
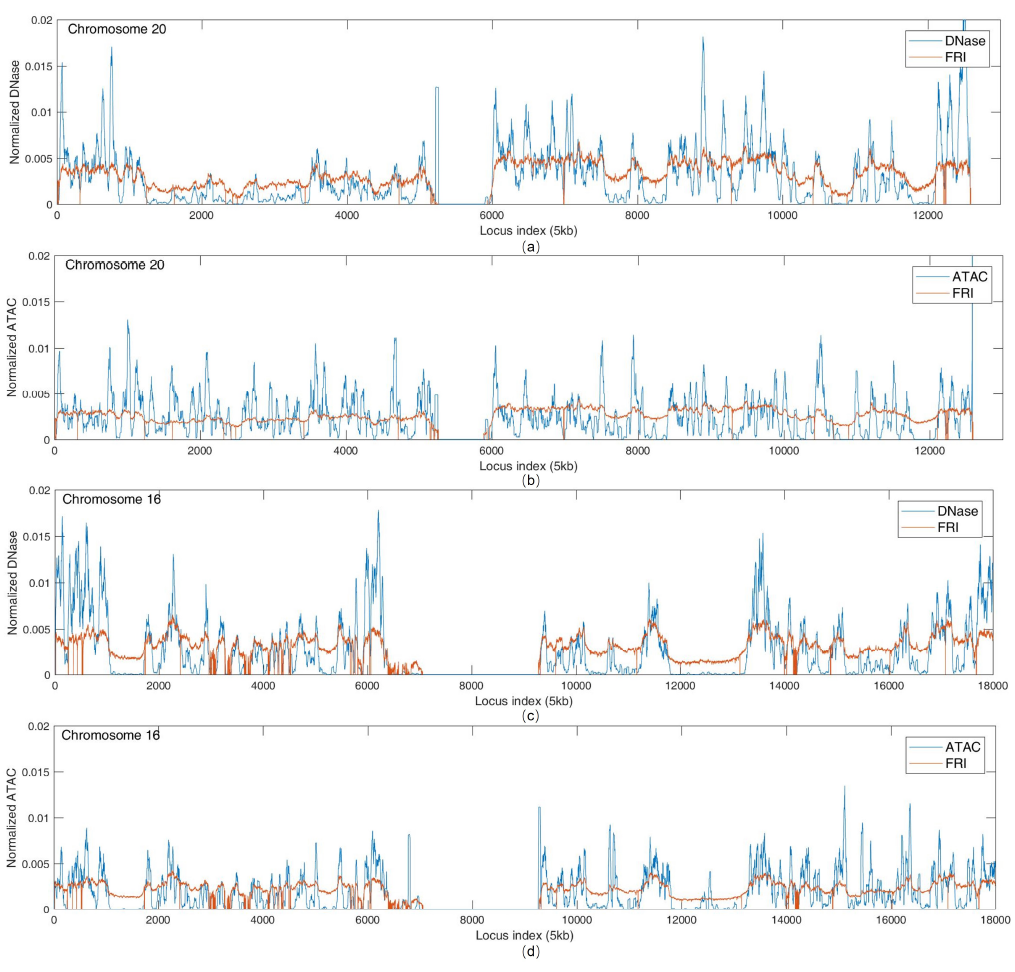
The illustration of the correlation between chromosome flexibility and chromatin accessibility. Two Hi-C data with resolution 5kb for chromosomes 20 (a, b) and 16 (c, d) of GM12878 are considered. The chromosome flexibility (red line) is calculated from our FRI model. The chromatin accessibility (blue color) is measured by DNase-seq (a, c) and ATAC-seq (b, d).

**Fig. 2.**
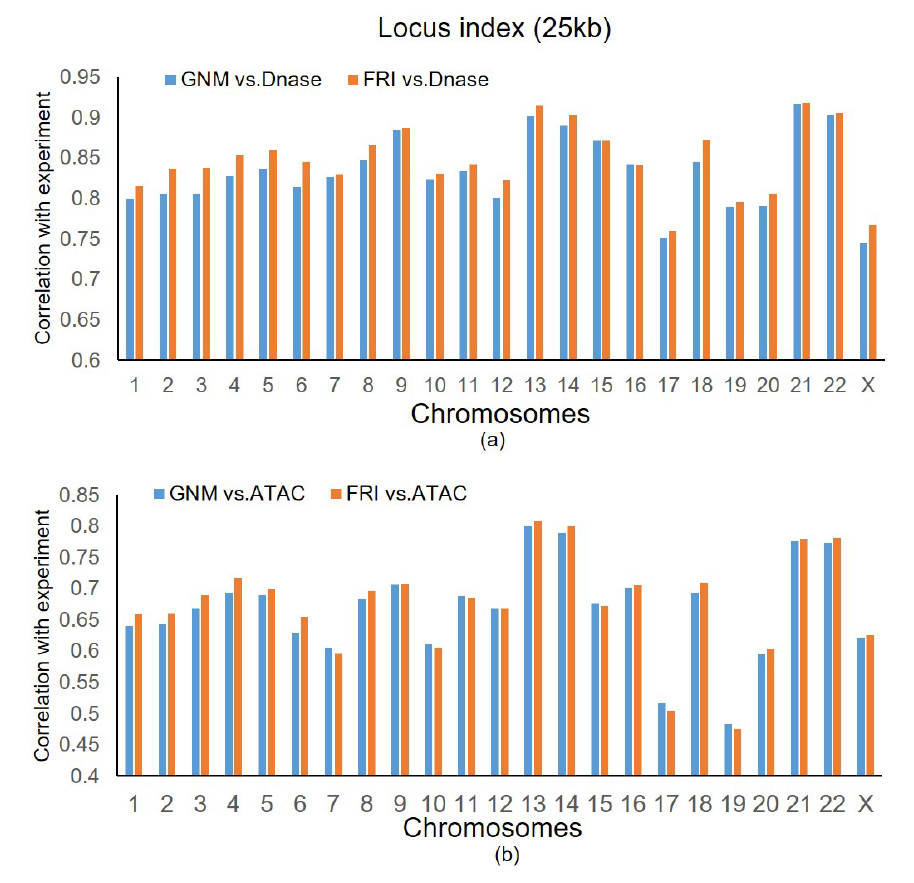
SCCs between chromosome flexibility and chromatin accessibility for GM12878 cell line. (a) SCCs of FRI (orange bar) and GNM (blue bar) for DNase-seq data. (b) SCCs of FRI (orange bar) and GNM (blue bar) for ATAC-seq data.

**Fig. 3.**
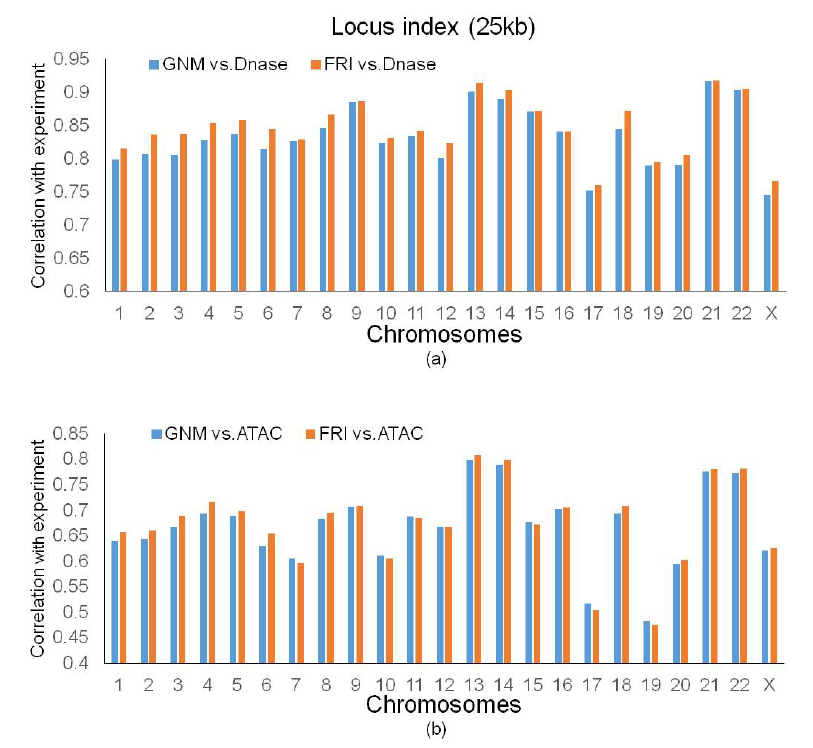
SCCs between chromosome flexibility and chromatin accessibility for IMR90 cell line. (a) SCCs of FRI (orange bar) and GNM (blue bar) for DNase-seq data. (b) SCCs of FRI (orange bar) and GNM (blue bar) for ATAC-seq data.

Currently, GNM is the only method that has been used to predict the chromatin accessibility from Hi-C data (Sauerwald *et al.*, 2017), to the best of our knowledge. In GNM, a mobility profile is generated from each chromosome. It is found that the mobility profile is linearly related to chromatin accessibility characterized by DNase-seq and ATAC-seq data. Larger mobility profile values indicate higher accessibility, whereas smaller values are associated with lower accessibility (Sauerwald *et al.*, 2017). Essentially, the mobility profile from GNM is similar to flexibility index in our FRI. To compare the performance of FRI and GNM in chromatin accessibility analysis, we calculate the SCCs for all 23 chromosomes in GM12878 and IMR90 using the GNM codes (Sauerwald *et al.*, 2017). The corresponding results from GNM are listed in Figures 2 and 3. It can be observed that, in nearly all the chromosome cases, SCCs from our FRI are higher than those from GNM in both DNase-seq and ATAC-seq. More specifically, in DNase-seq models, the average GNM (FRI) SCCs of all 23 chromosomes for GM12878 and IMR90 are 0.838 (0.860) and 0.833 (0.847), respectively. In ATAC-seq models, the average GNM (FRI) SCCs for GM12878 and IMR90 are 0.618 (0.633) and 0.667 (0.674). We can see that the average SCCs of FRI method are around 1% to 2% higher than GNM method in both DNase-seq and ATAC-seq.

### 3.3 Robustness of FRI for chromosome flexibility analysis

In the above section, we only consider the Hi-C data with resolution of 25kb. To further test robustness of FRI method, we consider the Hi-C data from the GM12878 cell line in different resolutions, i.e., 50kb and 100kb. Correspondingly, scale parameter *η* is set as 2.0 * 10^*−*4^ and 1.0 * 10^*−*4^, respectively. The SCCs are calculated for both ATAC-seq and DNase-seq data. The results are illustrated in Figure 4(a) and Figure 4(b). The GNM results are also listed for comparison. Consistent with the above results, our predictions are highly accurate and is constantly better than GNM. Interestingly, the accuracy of both models increases with the resolution.

**Fig. 4.**
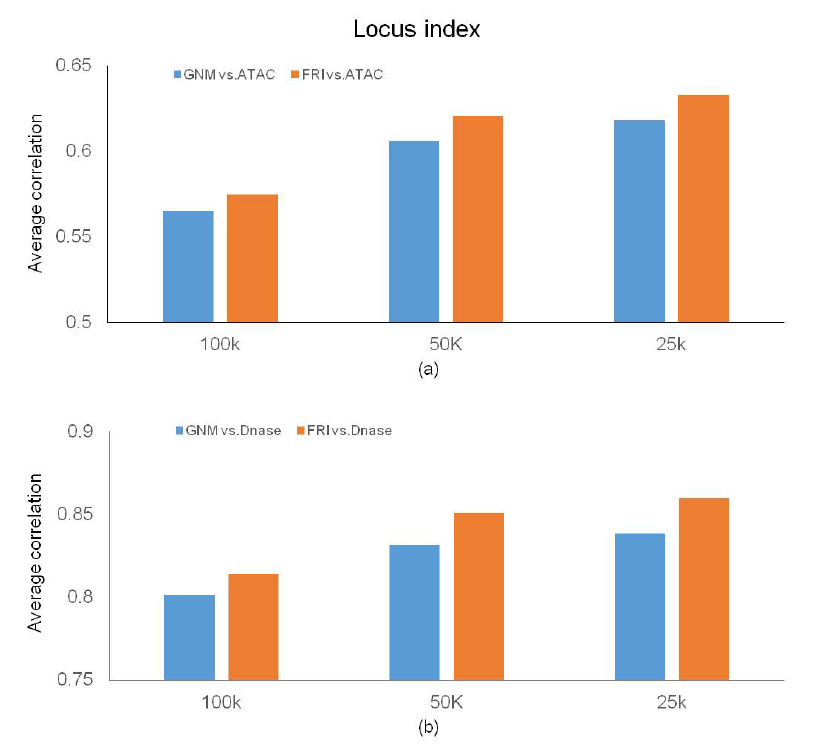
Average SCCs between chromosome flexibility and chromatin accessibility for all chromosomes from GM12878 cell line. (a) Average SCCs of FRI (orange bar) and GNM (blue bar) for ATAC-seq data. (b) Average SCCs of FRI (orange bar) and GNM (blue bar) for DNase-seq data.

### 3.4 Effect of inter-chromosome interactions on chromosome flexibility

Both intra-chromosome and inter-chromosome interactions have a great impact on the chromosome packing density, thus directly influence chromosome flexibility. However, the consideration of the inter-chromosome information will dramatically increase the computational costs, especially when matrix operations are involved. Therefore, it is prohibitively expensive for GNM to incorporate the effect of inter-chromosome interactions (Sauerwald *et al.*, 2017). Since FRI method involves only simple algebraic operations, it brings great promise to tackle the challenge from inter-chromosome interactions.

The Hi-C data from GM12878 with resolution 100kb is used. These are the highest resolution data with inter-chromosome interactions that we can obtain. Figure 5 demonstrates the SCCs between chromosome flexibility and chromatin accessibility for DNase-seq (a) and ATAC-seq (b). Two situations are considered for comparison. One is with only intra-chromosome interactions and the other is with both intra and inter-chromosome interactions. It can be seen clearly that the incorporate of the inter-chromosome information can further increase the accuracy of our chromosome flexibility model. For DNase-seq, the average SCC is 0.832 for both intra and inter case and it is 0.02 higher than the intra model (0.831). For ATAC-seq, the average SCC is 0.587 for both intra and inter case and it is 0.01 higher than the intra model (0.586).

**Fig. 5.**
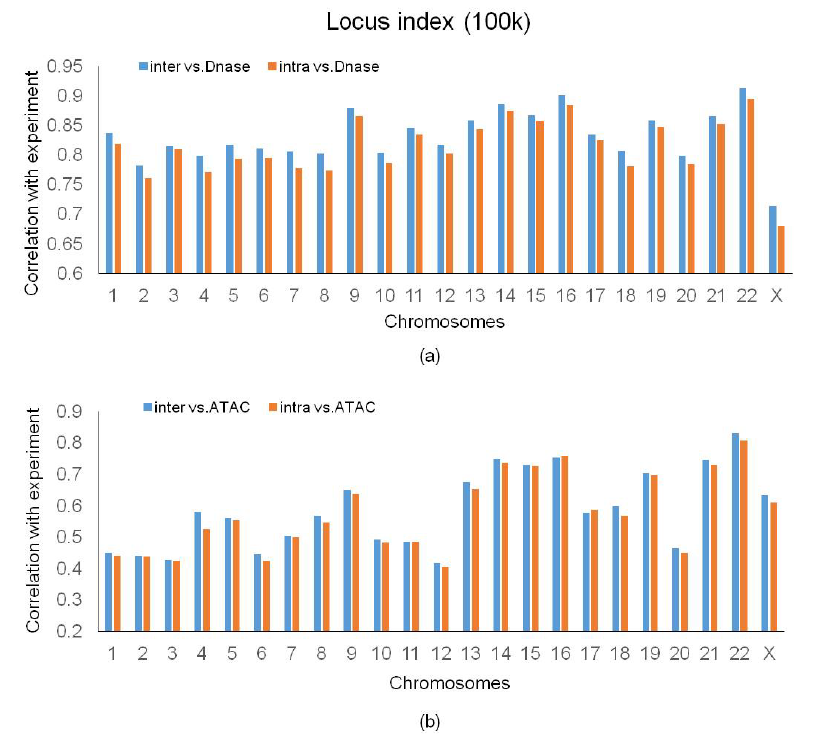
SCCs of FRI with inter-chromosome interactions for GM12878 cell line. (a) SCCs of FRI (orange bar) and GNM (blue bar) for DNase-seq data. (b) SCCs of FRI (orange bar) and GNM (blue bar) for ATAC-seq data.

### 3.5 Algorithm efficiency comparison between FRI and GNM on chromosome flexibility analysis

A significant advantage of FRI over all previous models in flexility analysis is its great efficiency and low computational cost. To compare the algorithm efficiency for FRI and GNM in chromosome flexibility analysis, we measure running times for all 23 chromosomes from GM12878 cell line. Both FRI and GNM are implemented with MATLAB. A linux server with Xeon(R) E5-2690 CPU (2.60GHz) and 512 GB memory is used.

Different resolutions (5kb, 25kb, 50kb, 100kb and 250kb) are considered and results are listed in Figure 6. Here the values represent the total computational time for all 23 chromosomes together. It can be seen clearly that FRI method is much more efficient than GNM method. For all resolutions, the running time of FRI is significantly less than that of GNM. Previously in GNM, it takes about 1.5 hour per chromosome at 5kb resolution using 10 CPUs (Sauerwald *et al.*, 2017). In contrast, it only costs FRI several minutes on a single CPU. These results are consistent with the model complexity (Xia *et al.*, 2013; Opron *et al.*, 2014). Essentially, FRI uses only simply algebraic operations, whereas GNM requires not only matrix operations but also eigenvalue decomposition.

**Fig. 6.**
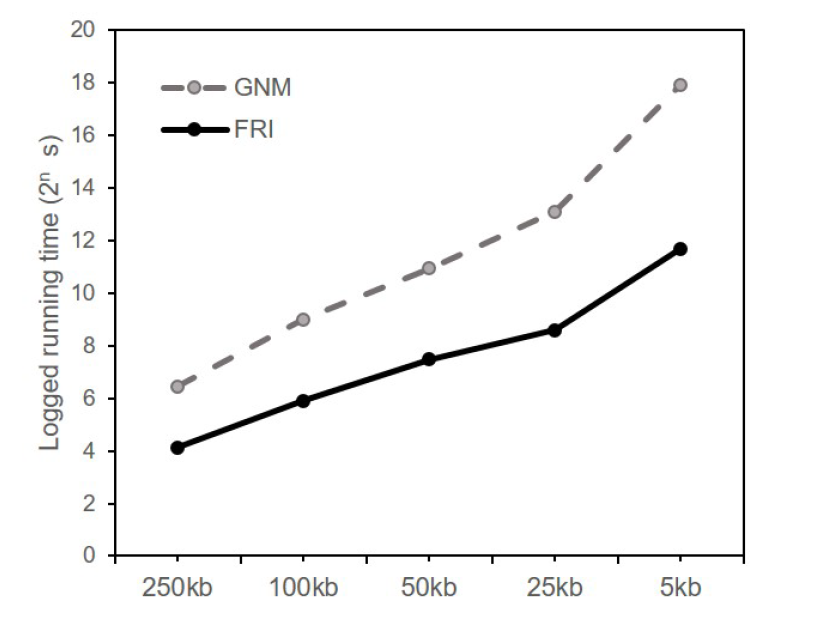
Running time of GNM and FRI on 23 chromosomes. The Hi-C data in different resolutions are considered. A significant reduce of the computational time in FRI can be clearly observed.

## 4 Conclusion

In this paper, the flexibility and rigidity index (FRI) model is introduced for the first time to analyze the chromosome packing, flexibility and dynamics. We evaluate the flexibility index for each locus. It is found that the flexibility index can be used to characterize the chromosome flexibility. A high correlation is found between our chromosome flexibility and accessibility measurements, including DNase and ATAC. Compared with the Gaussian network model (GNM), FRI is not only more accurate, but also significantly more efficient in both computational times and costs. Moreover, FRI can incorporate the inter-chromosome information into the flexibility evaluation, thus further enhance the model accuracy.

## Acknowledgements

The author Kelin Xia would like to thank Amartya Sanyal for his introduction and discussion of Hi-C experiments and data analysis.

